# Changes in spatial self-consciousness elicit grid cell-like representation in entorhinal cortex

**DOI:** 10.1101/2023.07.21.550007

**Authors:** Hyuk-June Moon, Louis Albert, Emanuela De Falco, Corentin Tasu, Baptiste Gauthier, Hyeong-Dong Park, Olaf Blanke

## Abstract

Grid cells in entorhinal cortex (EC) encode an individual’s location in space and rely on environmental cues and multisensory bodily cues. Body-derived signals are also primary signals for the sense of self as the continuous application of visuo-tactile bodily stimuli elicits illusory drifts in perceived self-location. It is unknown whether illusory changes in self-location are sufficient to elicit grid cell like representation (GCLR) in EC and how this compares to GCLR during conventional virtual navigation. Our results show that illusory changes in perceived self-location (independent of changes in environmental navigation cues and explicit imagined navigation) evoke entorhinal GCLR, correlating in strength with the magnitude of perceived self-location, and characterized by similar grid orientation as during conventional virtual navigation. These data demonstrate that the same grid-like representation is recruited when navigating based on environmental, mainly visual cues, or when experiencing illusory forward drifts in self-location, driven by perceptual multisensory bodily cues.

## Introduction

To localize oneself in space is a fundamental ability of many animal species, including humans. In the mammalian brain, several classes of neurons have been identified that encode information about an individual’s location and orientation in space. Grid cells in entorhinal cortex (EC), first described in rodents, represent a remarkable example of neurons that fire when the individual is at specific locations in space, and their firing fields form hexagonal grids tessellating the environment^1, 2^. Due to these characteristics, grid cells are thought not only to encode an individual’s location in space, but also to provide navigation vectors (i.e. displacement and direction) based on the periodicity of their grids^3, 4^. Grid cell-like modulation of brain activity has also been observed in humans using functional Magnetic Resonance Imaging (fMRI) as well as single-unit recordings^2, 5^. The hexadirectional modulation of fMRI signals observed in EC (grid cell-like representation; GCLR) arguably reflects the activity of grid cell populations^5–7^. While most of human grid cell studies have relied on environmental (mostly visual) cues during virtual navigation, studies in rodents have shown that grid cells integrate not only environmental cues but also sensory signals from the body (such as vestibular or somatosensory signals) to update self-location^1, 8, 9^. The dearth of data on the role of bodily signals in human grid cell studies is not surprising, given that in those studies the body of participants is required to remain stationary (either due to neurosurgically implanted depth-electrodes or constrained in the MRI scanner; for discussion and comparison with animal work see^10, 11^). While seminal human studies have described hexagonal grid codes also in navigation-related cognitive top-down processes (e.g., “navigation” in conceptual space, imagined spatial navigation), self-location in traditional human navigation studies, still mainly relies on *visuospatial cognitive* mechanisms, giving rise to a disconnect between the participant’s body ‘in’ the real world and the agent of virtual navigation ‘in’ the visual environment (for discussion see^11^). This renders the investigation of the role of *bodily perceptual* signals and their integration with visual environmental signals in spatial navigation research difficult. Recent research in human neuroscience on bodily self-consciousness^12–15^, however, developed several experimental methods that make it possible to investigate the role of bodily signals and their integration with environmental signals in human spatial navigation studies using fMRI^16^. Moreover, research on bodily self-consciousness demonstrated that bodily signals provide important sensory input to self-location^17–19^.

Under normal conditions, the self is experienced at the physical place occupied by one’s physical body and it has been shown that this co-location of self and body (i.e., embodied self) is based on the continuous integration of multisensory bodily signals (e.g., proprioceptive, tactile, motor, and visual-bodily cues)^12–15^. Yet, the locations of self and body do not always overlap and in certain neurological conditions the embodied self may be strongly altered: the self can be experienced at a different position than the individual’s body, for example as spatially dissociated from the individual’s body at an extracorporeal location, as in so-called out-of-body experiences^20, 21^. Subsequent research in healthy participants has demonstrated that comparable changes of the embodied self, characterized by extracorporeal self-location, can also be experimentally induced by combining conflicting multisensory stimulation and immersive virtual reality (VR)^12, 18, 22, 23^. These studies corroborate the idea that self-location, or the perceived spatial location of the self (whether at the location of the body or elsewhere in space), is an active and continuous brain process based on the integration of multisensory neural signals representing the individual’s body. For example, during the full-body illusion (FBI) paradigm^22, 24^, participants watch the back of an avatar (or the image of their own body) placed in front of them within a virtual environment, while receiving tactile stimulation to their back. During conditions eliciting a dissociation of perceived self-location from the individual’s own body, participants view the avatar being touched synchronously with the touches they receive on their back: prolonged exposure to such spatially conflicting multisensory stimuli (i.e., a synchronous signal in two spatially distinct locations, one for avatar, one for participant’s body; Fig.1a,b) is associated with participants reporting a forward drift in self-location towards the distant avatar (Fig.1a,b)^18, 22, 24, 25^ (see also^12, 26^). Thus, in these experiments, participants perceive self-location that does not overlap with their physical body position and perspective^17, 18^. It is currently not known whether such subjective and *passive* drifts in self-location, based on bodily perceptual signals (i.e., visuo-tactile stimulation) and independent of any virtual navigation, explicit imagined navigation, or the related visual environmental changes, are sufficient to activate entorhinal GCLR signals and how they compare to GCLR signals during *active* virtual navigation.

Here, we linked the body of participants in the MRI scanner with an avatar placed within a virtual environment through visuo-tactile stimulations (i.e., FBI) and investigated whether experimentally induced spatial dissociations, between the perceived location of the self and the physical location of one’s own body, would impact grid cell representation in human EC. More specifically, we hypothesized that experimentally-induced forward drifts in perceived self-location, an internal subjective change as elicited by the FBI, would be sufficient to elicit grid cell-related activity, without any overt virtual navigation. To test this hypothesis, we measured the hexadirectional modulation of fMRI signals in human EC (i.e., GCLR)^5, 7, 16^ during the induction of illusory changes in self-location for many orientation angles across an entire VR room, as elicited by periods of visuo-tactile stimulation in the FBI paradigm. We induced the FBI repeatedly from the same central spatial origin and for an avatar shown at the same distance in the virtual room (see Fig.1a-c). Critically, the avatar was shown at many different circular positions covering the entire virtual room, i.e., every 20° (Fig.1c). This allowed us to determine: (1) whether GCLR in EC can be induced by visuo-tactile stimulation and whether it depends on the synchrony of multisensory stimulation (lower GCLR for control asynchronous visuo-tactile stimulation), (2) whether such GCLR is related to the magnitude of perceived self-location change, and (3) how the entorhinal grid code, elicited by the FBI-induced changes in self-location, compares with the one associated to conventional virtual navigation. During the present FBI paradigm, the participant’s body, the avatar, and the visual environment all remained static, differing from previous virtual navigation fMRI studies^5, 7, 27^. The present procedure also differs from imagined or simulated virtual navigation^28, 29^, which taps into imagery-based top-down processes of *spatial cognition*^30^ as opposed to bottom-up processes of *bodily perception* as tested in the present FBI approach^15^.

Our results show that illusory changes in perceived self-location, systemically induced by FBI toward many circular directions across the virtual room, are sufficient to evoke entorhinal GCLR. This effect was only found in the main experimental condition inducing a stronger FBI associated with larger drifts in self-location (synchronous visuo-tactile stimulation), but not in the control condition (asynchronous stimulation), and the degree of the entorhinal GCLR response correlated with the magnitude of the change in perceived self-location. These data demonstrate grid cell-related activity in human EC, when our participants perceived a location of their bodily self that did not overlap with their physical body position, independent of changes in environmental visual cues (i.e., of first-person viewpoint or visual landmarks that are present in human grid-cell research during virtual navigation), and without recruiting top-down cognitive mechanisms involved in imagined or simulated virtual navigation^28, 29^. Additional data show that the grid orientation of the GCLR signal during FBI periods is similar to the grid orientation during periods of standard virtual navigation (button-press controlled virtual navigation movements in the same environment), indicating that humans recruit the same grid-like representation when navigating based on environmental, mainly visual, cues or when perceiving illusory forward drifts in self-location, driven by perceptual visuo-tactile cues.

## Results

### Drifts in self-location towards the avatar shown at many circular positions in the virtual room

Participants were exposed to a full-body illusion (FBI) paradigm with a novel and MRI-compatible design, adapted to the supine position required by MRI (Fig. 1; Methods; Supplementary Videos 1-3). To induce the FBI, we applied synchronous visuo-tactile stimulation that associated tactile stimulation of the participant’s body (i.e., abdomen) with touches that were seen as applied on the same body part of the virtual avatar (SYNC stimulation, Fig.1). A second multisensory condition (ASYNC stimulation) served as a control. During ASYNC stimulation visuo-tactile stimulation was applied asynchronously with a delay between felt and seen touches^24, 26, 31^. In order to test whether the FBI and the associated subjective drifts in self-location are sufficient to elicit GCLR activity (i.e., hexadirectional fMRI signal modulation), we designed the following experiment. We induced the FBI from the same central spatial origin (Fig.1c) and with an avatar shown at a distance of 2.5 virtual meters (vm; 1 vm in the VR environment was implemented to match 1 m in the real world) in the virtual room. Critically, the avatar was shown at 18 different circular positions in the virtual room, with 20° resolution and covering the entire virtual room (360°/20° = 18 positions; Fig.1c). Hence, visuo-tactile stimulation (in the SYNC or ASYNC condition) was carried out for each of these 18 circular positions during each FBI-induction session. After each visuo-tactile stimulation phase, the avatar disappeared and the viewpoint was shifted to a different position (backward shift to a random position; 2∼3 vm away from the central spatial origin) and participants indicated their perceived self-location by navigating to the position where they felt to be located during visuo-tactile stimulation (self-location report; Fig. 1g).

**Fig. 1.**
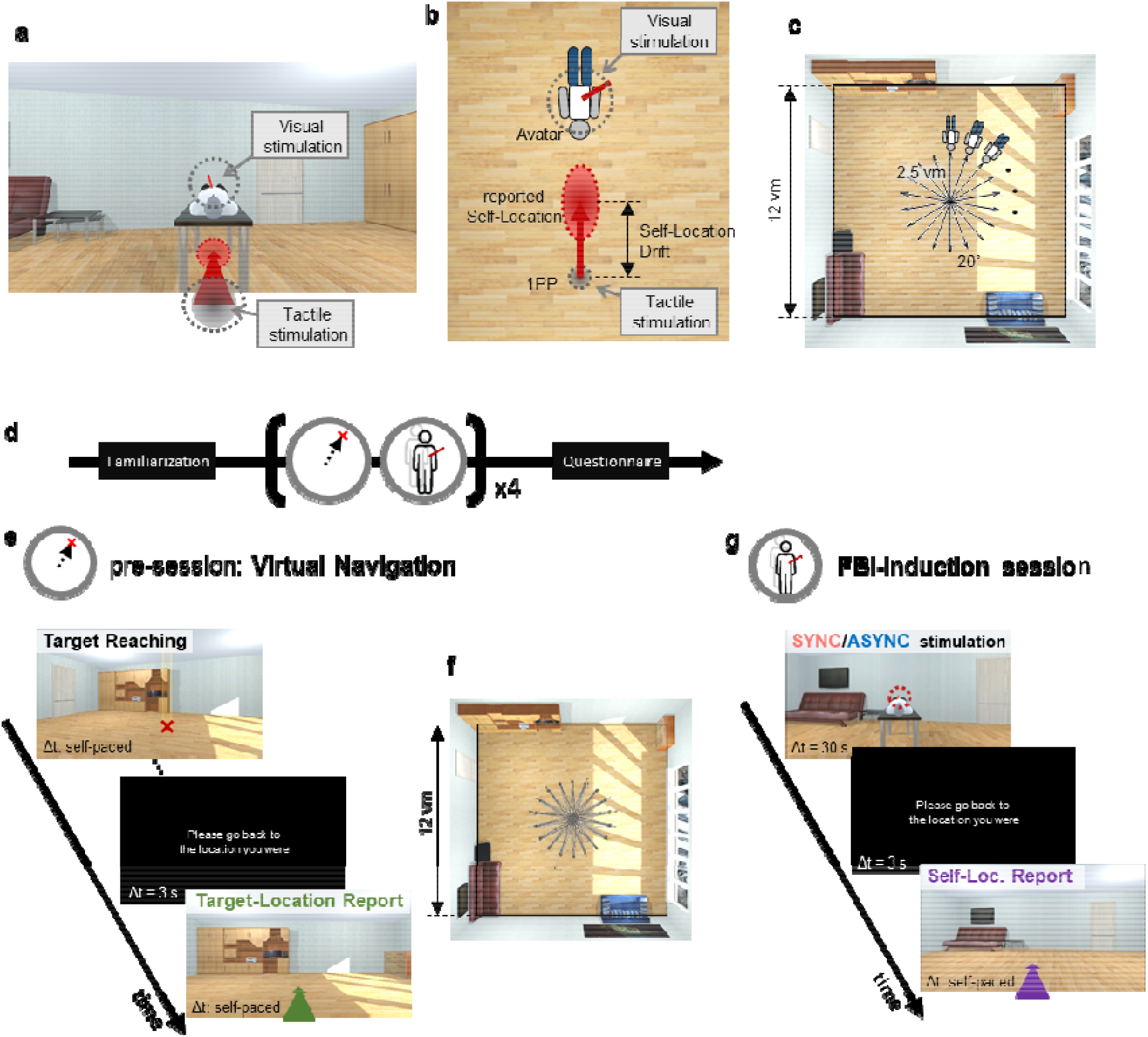
MRI-compatible full-body illusion (FBI) paradigm to estimate grid cell-like representation (GCLR) **a,** The first-person perspective (1PP) scene seen by our participants during the FBI paradigm in the MRI scanner. To induce FBI in the scanner, a virtual avatar in a supine position – thus congruent with the participant’s physical body position – was placed 2.5 virtual meters (vm) in front of the first-person viewpoint anchored at the center of the virtual room. Of note, the participant’s physical body was not visible nor shown in VR as in the conventional FBI with a standing avatar. Then, the abdomen of the participants and corresponding body parts of the avatar were stroked for 30 sec (i.e., visuo-tactile stimulation). **b,** Top view of the virtual environment shown in panel a. Visual stimulation on the distal avatar and tactile stimulation on participant’s physical body are associated during FBI. The association of visuo-tactile stimuli in two distinct locations induced drift in self-location from 1PP location towards the avatar. **c,** During the illusion-induction sessions, the FBI was induced toward every 20° direction spanning the virtual room (total of 18 trials per session), so that GCLR could be estimated. **d,** Participants went through our MRI paradigm composed of four pairs of virtual navigation pre-session and FBI-induction session. They were exposed to the familiarization procedure before the main task. A questionnaire was carried out at the end of the experiment. **e,** In the virtual navigation pre-session, participants performed simple target-reaching tasks, freely navigating to the target indicated by the yellow light pillar (also marked with the red ‘x’; which was not shown to the participants). At random trials, they were asked to report the self-location experienced during the preceding trial by actively navigating (target-location report; green). **f,** The virtual navigation pre-session was designed to match the presumed spatial vectors during the FBI-induction phase, providing comparable and robust GCLR data. The figure shows the navigation traces of an exemplary participant during the virtual navigation pre-sessions, which looks similar to panel c. **g,** FBI-induction session consisted of two phases: visuo-tactile (SYNC/ASYNC) stimulation and self-location report. During visuo-tactile stimulation in the SYNC condition, the full body illusion was induced by synchronous visuo-tactile stimulation associated with a virtual avatar for 30 sec. In the control stimulation condition (ASYNC), asynchronous stimulation between the participant and the avatar was applied. During self-location report phase following each visuo-tactile stimulation phase, participants were asked to report the self-location perceived during the stimulation in order to quantify the perceived self-location drifts. At the beginning of the self-report, they were placed behind the center of the room (between 2 and 3 vm) and, then, had to actively navigate to perceived locus of stimulation. (Also, see supplementary videos 1-3).

In line with previous studies about perceived self-location^18, 22, 24, 32^, during the self-location report, participants indicated a forward drift in self-location towards the avatar induced by the visuo-tactile stimulation in the MRI scanner. The drift was significantly larger in SYNC than ASYNC condition (mixed-effects regression; df = 1, F = 43.8, d = 0.28, p = 4.94e-11, n = 32; Fig. 2a). We performed an additional drift analysis that corrected the reported self-location (after SYNC and ASYNC trials) with the reported self-location during the preceding virtual navigation pre-session. This analysis again confirmed a larger drift in self-location towards the avatar in the SYNC vs. ASYNC condition (df = 1, F = 1629.2, d = 0.37, p < 2.2e-16, n = 32; Supplementary Fig. 2c-d). We also verified that the different viewing angles in the room could not account for the observed difference in reported self-location between the conditions through Two-way repeated measures ANOVA (Df = 17, F = 0.301, p = 0.997; Supplementary Fig. 3).

**Fig. 2.**
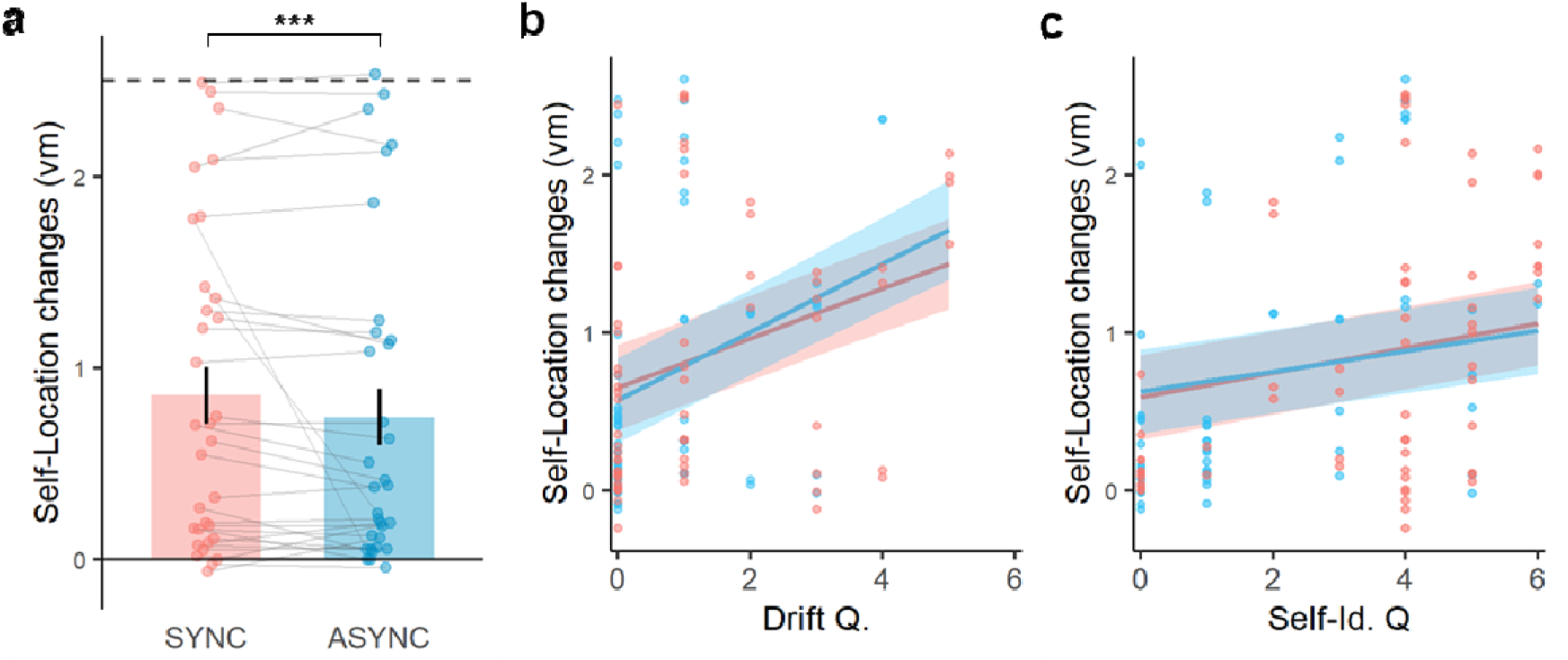
Visuo-tactile stimulation induces forward drift in perceived self-location in the environment. **a,** The perceived self-locations reported following the stimulation phase were farther away from the viewpoint (and closer to the avatar location) in the SYNC condition compared to the ASYNC condition (p = 4.94e-11; n = 32). Here, 0 vm corresponds to the first-person viewpoint of the participants during the illusion phase (i.e. center of the room), while the dashed line at 2.5 indicates where the virtual avatar was placed. **b,c,** The higher questionnaire ratings of both Drift Question (b) and Self-identification Question (c) were associated with the greater self-location changes (Drift Q: p < 2.2e-16, n =32 / Self-Id. Q: p = 3.45e-16, n = 32). The linear relations between the two variables were separately estimated using linear mixed-effects models with the trial-wise data points. Each dot in the plots represents the condition-wise mean value of Self-Location change for each participant for visualization. The error bars indicate standard errors of the mean. *** p < 0.001.

To further ensure the successful induction of the FBI, we also administered a questionnaire, at the end of the experiment (Supplementary Fig. 1, see Methods). Participants reported stronger subjective forward drifts in self-location (Drift Q.; r = 0.42, p = 0.017, n = 32) in the SYNC condition than in the ASYNC condition. In addition, illusory self-touch for seen touches applied to the avatar (r = 0.81, p = 4.67e-06, n = 32), self-identification with the avatar (Self-Identification Q.; r = 0.62, p = 4.56e-04), and the feeling of presence (Presence Q.; feeling of being within the VR environment; r = 0.49, p = 5.4e-3) were also significantly higher in the SYNC than ASYNC condition. No effect between conditions was found for the control question (r = 0.29; p = 0.11). Additional correlation analysis revealed that participants’ drift in perceived self-location (behavioral response measured by active virtual navigation) was correlated with the subjective feeling of forward drift in self-location (on questionnaire ratings reported at the end of the experiment; Drift Q.; F = 142.5, *pη*^2^ *= 0.08,* p = 2.2e-16, n = 32; Fig. 2b) as well as self-identification (df = 1, F = 67.9, *pη*^2^ *= 0.04,* p = 3.45e-16, n = 32; Fig. 2c). The other questionnaire ratings (i.e., self-touch and presence Q.) were not correlated with drift in self-location. These findings show the successful translation of an FBI paradigm to the supine position as well as its integration into a spatial navigation setup in the MRI scanner. Critically, applying visuo-tactile stimulation, we successfully induced forward drifts in perceived self-location within the virtual room that were directed away from the center of the virtual room and extended along 18 different circular directions, enabling us to investigate its impact on GCLR.

### GCLR and drift in perceived self-location within the virtual room are linked

Before evaluating the GCLR during visuo-tactile stimulation, we assessed GCLR in the virtual navigation pre-sessions during which participants were actively navigating in the VR room (Methods; Fig. 1e). These pre-sessions were designed both to assess the general presence of entorhinal GCLR in our participants using already established methods (i.e., virtual navigation^5, 7, 16, 27^), and to robustly estimate the grid orientation for each participant, which could then be used to investigate GCLR during the visuo-tactile stimulation phase (i.e. FBI). We calculated GCLRs separately for the target reaching and target-location phases, associated with different cognitive processes. Our results show significant GCLR during the target-location report phase (r = 0.45, p = 5.57e-03, n = 32), but not during the target reaching phase (r = 0.07, p = 0.34, n = 32; Supplementary Fig. 4; further discussed in the Supplementary Text). The significant GCLR during the target-location report phase shows that a series of short and discrete navigations in an indoor virtual room, which were expected to be comparable to the FBI-induced self-location changes in our study, are sufficient to evoke GCLR in the human EC.

Next, we assessed whether the induced illusory FBI states could generate grid cell-like activity in EC. We hypothesized that, if illusory drifts in self-location across the virtual room recruit the identical (or similar) grid code, active during virtual navigation in the same virtual space, then the GCLRs from the different experimental phases (e.g., FBI-induction phase and active virtual navigation phase) should maintain the grid orientation^28^. On this premise, we calculated GCLRs during the three different phases of the FBI experiment (i.e., SYNC stimulation phase, ASYNC stimulation phase, self-location report phase), as based on the grid orientation estimated from the virtual navigation pre-sessions (i.e., the target-location report phase; Fig. 1). This analysis revealed a significant GCLR activation during the FBI-inducing SYNC condition (r = 0.35, p = 0.024, n = 32; Fig. 3b-red), but not during the ASYNC condition (r = 0.20, p = 0.127, n = 32; Fig. 3b-blue). This shows that specific changes in perceived self-location - that were induced for the avatar shown at a distance, in different trials across many orientations in the virtual room, and without any ongoing virtual navigation - activated entorhinal GCLR. For the subsequent self-location report phase (participants virtually navigated to indicate their self-location during the immediately previous FBI-induction session), we again detected a significant GCLR activation (also based on the grid orientation estimated from the pre-sessions; r = 0.47, p = 3.67e-03, n = 32; Fig. 3b-purple). Moreover, the rotational symmetry of the heading-direction-dependent BOLD signal was found to be significant only for the six-fold (i.e., GCLR) symmetries in the SYNC condition and the self-location report phase, but not for the other control symmetries (4,5,7-fold) and never in the ASYNC condition (Fig. 3d)

**Fig. 3.**
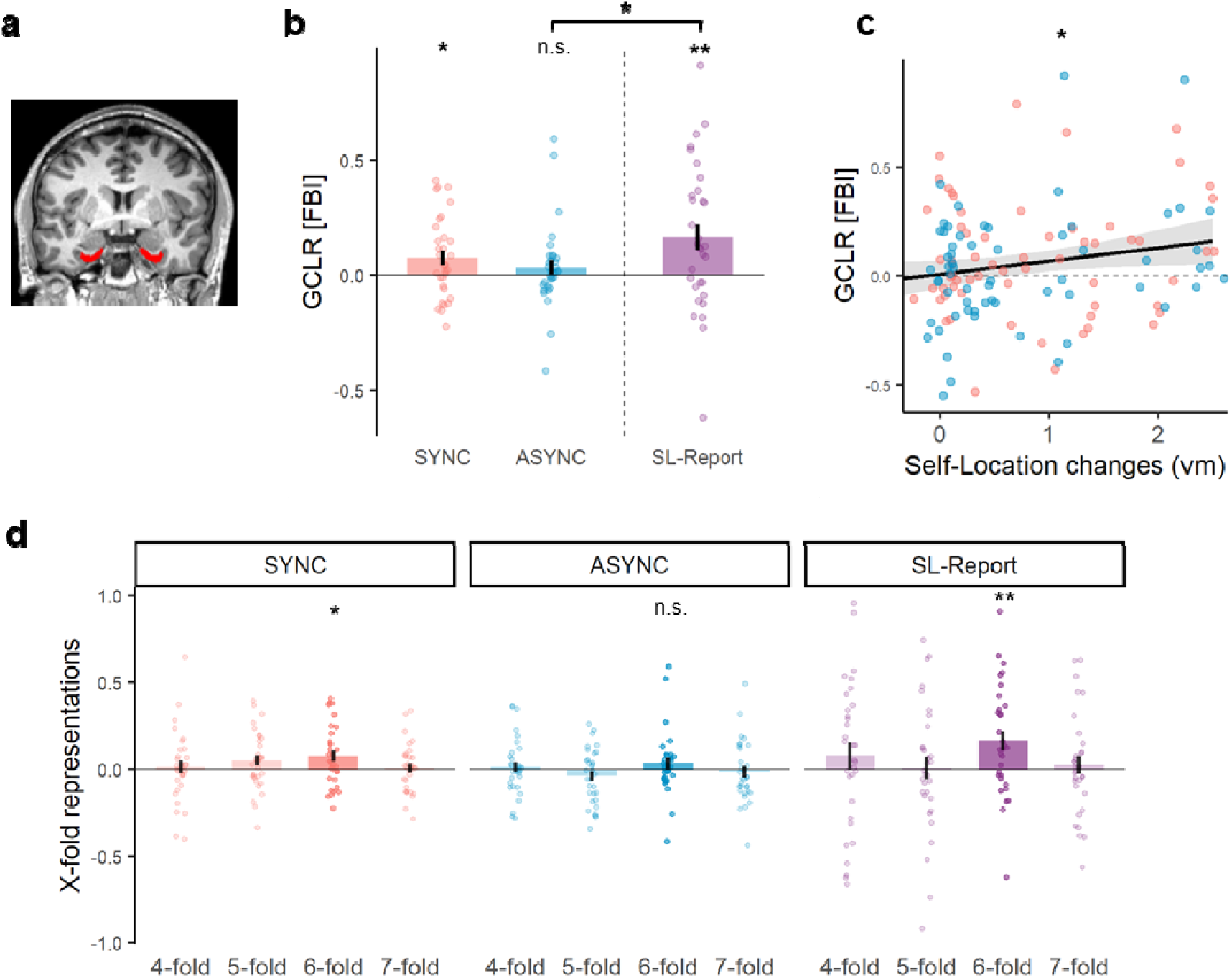
GCLR during the FBI was related to the drift in the perceived self-location and its grid orientations were similar to the virtual navigation. **a,** Bilateral entorhinal cortex was anatomically defined for each participant in the native space. **b,** GCLRs during the FBI-induction session were calculated based on the grid orientation estimated from the virtual navigation pre-sessions (see Supplementary Fig. 4). Significant GCLR was observed in the SYNC condition of the simulation phase (i.e. FBI), but not in the ASYNC condition. GCLR was also detected during the self-location report phase (purple), where the participant actively navigated to the self-location perceived during the preceding stimulation phase. The amplitudes of the GCLR generated during the self-location reports were greater than during the ASYNC stimulation phase, while no significant difference was found with the SYNC stimulation phase. Each error bar indicates the standard error of the mean. **c**, The amplitudes of the GCLR during the visuo-tactile stimulation were significantly correlated with the drift in the perceived self-location (p < 0.05). For panel c, the statistical assessment was performed with a linear mixed-effects model based on the session-wise data. **d**, Rotational symmetries for heading-direction dependent GCLR BOLD signal were not significant, except for ‘6-fold’, during any experimental phase in the FBI-induction session, confirming that our GCLR results were specific for the hexadirectional modulation (i.e., 6-fold). n.s.: p >= 0.05, * : 0.01 <= p < 0.05, ** : 0.001 <= p < 0.01.

### Drift in perceived self-location within the virtual room correlates with entorhinal GCLR amplitude

Next, we investigated whether the strength of GCLR during FBI induction was related to the magnitude of the illusory drift in self-location. For this purpose, we assessed the relationship between the amplitude of GCLR in EC and the magnitude of the reported forward drift in self-location, by using a mixed-effects model on the session-wise data. Congruent with our hypothesis, this analysis revealed that the two measures were significantly correlated: larger forward drifts in self-location were associated with a stronger GCLR activity in EC (F = 4.52, *pη*^2^ *= 0.12,* p < 0.041, Fig. 3c). We found that this was observed irrespective of whether stimulation was carried out in the SYNC or ASYNC conditions: Thus, drifts in self-location, but of different magnitudes, were induced in both conditions and covaried with GCLR (drifts and GCLR were larger in the SYNC condition; see Fig. 3c). Using mediation analysis, we further investigated the relationship between synchrony of visuo-tactile stimulation, drifts in self-location, and GCLRs during the stimulation phase. This analysis revealed that the GCLR modulation was mediated by the changes in the perceived self-location (p = 0.041; Supplementary Fig. 7b), rather than solely influenced by the synchrony of the visuo-tactile stimulation (p = 0.35).

### GCLR induced by illusory drift in self-location has a similar grid orientation to GCLR induced by virtual navigation

The significant GCLRs in the SYNC phase (Fig. 3b-red) and in the self-location report phase (Fig. 3b-purple) during the FBI-induction session (both calculated based on the grid orientation during virtual navigation pre-session) suggest that grid orientation is maintained across different sessions and phases of the experiment. To further investigate the GCLRs associated with the two distinct types of self-location change (i.e., forward movements during virtual navigation vs. FBI-induced forward drifts in self-location), we performed additional analyses of grid orientations. Independent estimations of the grid orientation during the three phases of the FBI-induction session (i.e., SYNC/ASYNC stimulation, and self-location report) revealed that the grid orientations during the visuo-tactile stimulation phases (both SYNC and ASYNC) were similar to the grid orientations during the self-location report phase (absolute circular mean difference between orientations < 15°; SYNC: 1.15° ± 4.82°, Z = 7.84, r = 0.350; ASYNC: 2.36° ± 6.75°, Z = 4.46, r = 0.264, n = 64; Fig. 4a). This effect suggests that self-location changes induced by both visuo-tactile stimulation and active virtual navigation are characterized by the same grid representation (and arguably same cognitive map). Of note, the comparison of the grid orientation was performed for each session, between the visuo-tactile stimulation phase (either SYNC or ASYNC) and paired self-location report phase (i.e., within-session). In addition, comparing the grid orientation during the virtual navigation pre-session with the grid orientation during the visuo-tactile stimulation phase (i.e., SYNC and ASYNC) independently estimated (Supplementary Fig. 6), we found again that the grid orientation during SYNC was similar to the one during the virtual navigation pre-session and differed by a mean value of 10.49° ± 7.11° (Z = 4.11, r = 0.253, n = 64). In contrast, the mean difference in grid orientation between ASYNC and the virtual navigation pre-session was not statistically clustered (Z = 0.29, r = 0.067, p = 0.75, n = 64) and its confidence intervals could not be estimated (mean: 53.24°). These results are congruent with the significant GCLR in the SYNC (Fig. 3b-blue) and the non-significant GCLR in the ASYNC condition (Fig. 3b-blue): the significant detection of GCLR largely depends on the stable estimation of the grid orientation^5, 33^, as also indicated by the relationship between the grid orientation difference and GCLR (Supplementary Fig. 5a). Notably, the subject-wise absolute grid orientations varied and were uniformly distributed across subjects (virtual navigation pre-session: Rayleigh’s Z = 0.31, r = 0.10, p = 0.74; SYNC: Z = 1.10, r = 0.19, p = 0.34; ASYNC: Z = 1.21, r = 0.19, p = 0.30; self-location report: Z = 0.68, r = 0.15, p = 0.51), excluding the possibility that the correlations are driven by a few extreme data points associated with specific visual-inputs or landmarks.

**Fig. 4.**
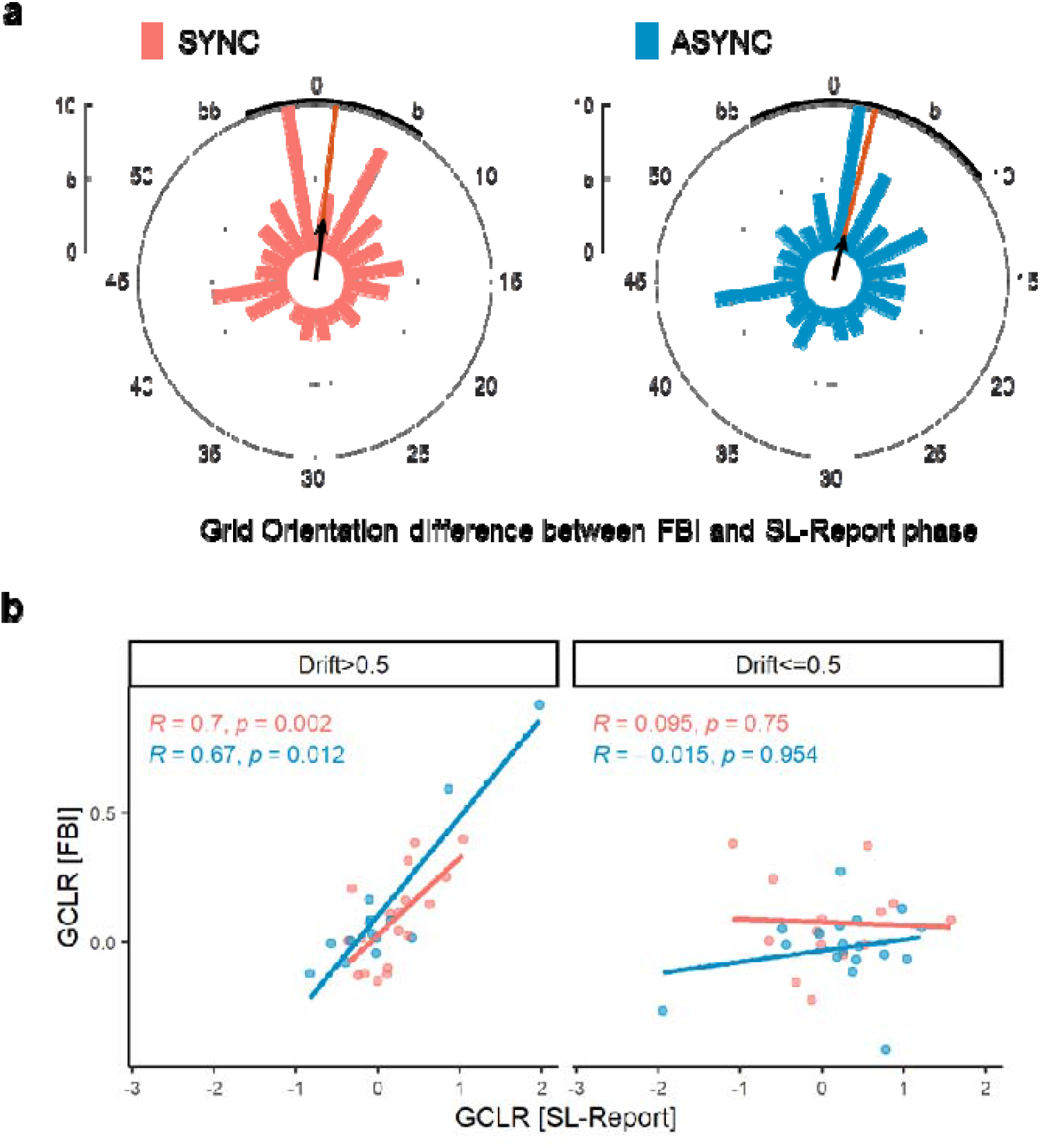
Grid codes for FBI-induced self-location drift and for standard virtual navigation during following self-location report are similar. **a,** The circular histograms show the distribution of angular differences between grid orientations of GCLR during the visuo-tactile stimulation phase and the immediately following self-location report phase. Grid orientation during the SYNC (red) and the ASYNC (blue) stimulation phases were similar to the grid orientation during self-location report phase. The ranges of the grid orientations and their angular difference are 0 to 60° for the GCLR of six-fold symmetry. A red line indicates the mean angular difference and the bold black curve at the rim indicates its confidence intervals. **b,** GCLR of individuals per stimulation condition (n = 32 * 2) were median-split into two groups based on the reported forward self-location drift (either Drift > 0.5 vm or Drift <= 0.5 vm), regardless of the conditions. Then, for each group and condition, we explored the relationship between GCLR during visuo-tactile stimulation phase and GCLR during the following self-location report phase. The results showed that when the drift was over 0.5 vm, GCLR during visuo-tactile stimulation phase was proportional to GCLR during the self-location report phase, regardless of the condition (SYNC or ASYNC). This suggests that similar grid cell-like activity was generated during the two phases with the matched spatial vectors. No significant relationship was observed when the drift was smaller than (or equal to) 0.5 vm, possibly due to the non-significant GCLR in these cases (see Supplementary Fig. 7a). Each correlation was assessed by the Spearman correlation.

Further supporting similar grid code during FBI and virtual navigation, we found that GCLR during the visuo-tactile stimulation phase is significantly correlated with the GCLR during the following self-location report phase (with matched spatial vectors; see methods). Critically, this was only found for larger illusory drifts (> 0.5 vm; SYNC: <u>ρ</u> *<u>= 0.70,</u>* p = 0.002, ASYNC: <u>ρ</u> *<u>= 0.67,</u>* p = 0.012) and absent for small drifts with an amplitude below 0.5 vm (Fig. 4b). This drift-dependent correlation again associates the GCLR during visuo-tactile stimulation with the illusory drift, while excluding the possibility that the correlation was merely derived from the timely intermingled fMRI dataset of the two phases. This finding also suggests that a minimum magnitude of self-location change is required to robustly estimate entorhinal GCLR and to reliably assess the similarity between GCLRs in different conditions (see Supplementary discussion; Supplementary Fig. 7a). These data provide firm evidence that the entorhinal GCLR during subjective forward drifts in self-location has a similar grid representation as compared to when participants actively navigate in the same virtual room.

## Discussion

We report that illusory changes in the perceived location of the bodily self, within a virtual environment, but without any virtual navigation, can elicit GCLR in human EC. We achieved this illusory perception of a direction-specific forward drift in self-location by multisensory bodily stimulation applied to participants while they were immersed in VR in an adapted FBI paradigm for MRI. Inducing the FBI repeatedly from the same central spatial position and along several angular directions covering the entire virtual room allowed us to detect entorhinal GCLR during the illusion-inducing synchronous visuo-tactile stimulation (SYNC stimulation phase of FBI-induction session; Fig. 3b). Such entorhinal GCLR was observed without any active virtual navigation and without any viewpoint changes that are generally used to elicit GCLR in humans. GCLR was absent in the asynchronous control condition that was matched in all aspects except for the delayed timing between visuo-tactile stimulations.

Using various paradigms, previous fMRI studies have investigated cognitive brain mechanisms of first-person visual perspective^34, 35^, recognition of visual environments^36, 37^, and mental imagery for spatial locations^38, 39^. Concerning grid-cell related brain activity in humans, spatial navigation research has investigated how entorhinal GCLR is modulated by changes in first-person perspective (i.e., virtual navigation)^5^, by mental imagery for spatial locations^29^, or by asking participants to imagine virtual navigation^28^. Here, we investigated whether perceptual bodily signals, applied during the FBI to manipulate the perceived spatial location of the bodily self (Fig. 2a) within a virtual room, would be sufficient to elicit and modulate entorhinal GCLR. Integrating previous self-location fMRI paradigms (e.g., ^12, 18, 22, 24, 26, 40^) and applying visuo-tactile stimulation between the abdomen of supine participants and a supine avatar shown at a distance in the virtual room, we induced drifts in self-location towards the distant avatar. Critically, such self-location drifts in the horizontal plane (comparable in magnitude to previous non-MRI VR studies^18^) were elicited in the MRI scanner along 18 circular directions spanning the entire virtual room, thereby enabling us to perform fMRI-based GCLR analyses (i.e., detection of hexadirectional fMRI signal changes). The present data show that GCLR in EC is activated by an illusory change in the perceived position of our participants’ self in the virtual room when they are passively exposed to visuo-tactile stimulation. This was achieved without changing any of the environmental visual cues that are tested in most previous human grid cell research, such as changes in viewpoint or visual landmarks during virtual navigation. Moreover, the observed GCLR in the SYNC condition was comparable in amplitude to GCLR during active navigation (Fig. 3b) and, compatible with previous work using virtual navigation^5, 16, 27^, showing a sixfold-specific signal modulation (Fig. 3d). These data extend previous work on entorhinal GCLR and the sense of self^16^. In contrast to the present study, Moon et al. (2022) did not apply visuo-tactile stimulation and did not show an avatar in a distant position, but in a spatially overlapping position with the participant’s body during an active virtual navigation paradigm. As the paradigm in the former study did not elicit systematic illusory drifts in self-location, it did not allow to investigate whether illusory drifts in perceived self-location are reflected in the GCLR signal, which was the main hypothesis of the present work. Drifts in self-location have been induced previously using multisensory stimulation in the MRI: Ionta et al. (2011)^24^ elicited drifts in the vertical plane associated with angular gyrus and posterior temporo-parietal activation, and Ehrsson and colleagues^26, 40^ measured changes in self-location by questionnaire responses (no drift measurements) associated with activation in retrosplenial cortex, intraparietal sulcus, posterior cingulate cortex, and hippocampus. Critically, these previous works did not investigate drifts in perceived self-location across many angles in a virtual environment and never with respect to GCLR in EC. The present data firmly link EC and virtual navigation with bodily self-consciousness (for review, Blanke et al., 2015^15^), in particular perceived self-location, by showing that illusory forward drifts towards the avatar’s body in the virtual room activate entorhinal GCLR, as if participants had actively navigated in the same virtual room. We note that to quantify these systematic changes in the location of the bodily self, we asked our participants to indicate their perceived self-location by actively navigating to the location where they had perceived their self in the virtual room during visuo-tactile stimulation. This was done immediately after each visuo-tactile stimulation phase to match the spatial vector (i.e., direction and goal of the displacement) of virtual navigation during the self-location report phase with the one during the preceding phase with illusory change in self-location. Thus, unlike previous human GCLR studies that used long and continuous virtual navigations (i.e., ∼ 80 vm long navigations within a diameter of 110 vm in the case of Moon et al., 2022; Doeller et al., 2010 used a circular arena with a diameter of 180 vm), and with participants freely changing their heading directions, the present experiment was designed to generate a series of short (maximally 2.5 vm by design) and discrete self-location changes while the heading direction was fixed during each FBI trial. By demonstrating that our virtual navigation control sessions - composed of similar navigation vectors (i.e., a series of short, discrete, and heading-direction-fixed vectors in the same virtual room) - could elicit significant GCLR, we argue that GCLR during synchronous visuo-tactile stimulation (SYNC condition) results from the drifts in perceived self-location.

The present GCLR effects are specific, because they were absent in the ASYNC control condition that elicited significantly shorter drifts in self-location. The ASYNC control condition fully matched both, visual and tactile, inputs of the SYNC condition and only differed in the timing of the applied visuo-tactile stimulation, rejecting the possibility that the GCLR we observed in the SYNC condition resulted merely from unimodal sensory inputs (i.e., either the visual or the tactile information alone) or from multisensory visuo-tactile stimulation per se (both SYNC and ASYNC were multisensory). We also note that the absolute grid orientations in the room varied across subjects and were uniformly distributed, further ruling out the possibility of visual input-driven modulation of EC activity. It could be argued that visual differences, related to the heading directions, generated the hexadirectional fMRI pattern (i.e., different viewing angles related to certain visual landmarks in the virtual room), yet, the reported GCLR cannot be based on such visual differences, as visual inputs were fixed (i.e. participants were oriented in the same directions) in both SYNC and ASYNC conditions.

One may argue that other factors (e.g., social or attentional components) could contribute to the GCLR in SYNC condition. For instance, GCLR might arise from encoding the avatar’s location (similarly to the GCLR during observed navigation^41^). However, we believe this is unlikely because the “social” avatar in the room was stationary (fMRI-based GCLR has not been detected for encoding any stationary agent). Even if it existed, it should not differ between SYNC and ASYNC conditions. Also, we believe that contributions of attentional components to our GCLR results are not very likely because tactile stimulation, visual stimulation, as well as multisensory visuo-tactile stimulation were all the same in both conditions and only the timing between visual and tactile stimulation differed between both conditions. Moreover, the observed relationship between GCLR and magnitude of self-location changes (Fig. 3c) as well as other drift-dependent GCLR results (Fig. 4b & Supplementary Figure 7a) cannot be explained by any of these accounts. We, therefore, argue that the observed differences in GCLR stem from the illusory self-location changes induced by multisensory stimulation and related processes. We cannot entirely exclude the possibility that the vividness (or strength) of the subjective experience of forward drift contributed to the difference in GCLR, because it was correlated with the magnitude of actual drift (Fig. 2b). Critically, we believe that this does not undermine the overall conclusions of our study. Even if potential other subjective experiences may have played a role (and especially as we explicitly induced subjective changes across conditions), it is still compatible with our study hypothesis and main conclusion that subjective changes in bodily self-consciousness as induced by multisensory stimulation elicits GCLR.

We do not think that earlier GCLR work by Horner and colleagues^28^ is directly related to the present data. These authors reported that the grid orientation between imagined virtual navigation and standard virtual navigation was similar, suggesting comparable coding for entorhinal GCLR during imagined virtual navigation and standard virtual navigation (i.e., two spatial cognitive processes). Inspired by these findings, we hypothesized that the illusory forward drifts in perceived self-location (as induced by multisensory stimulation and related perceptual processes) would recruit a similar grid code (with the same grid orientation) as virtual navigation in the same virtual room (pre-session) and, therefore, also estimated GCLR during the FBI, but now based on the grid orientation as defined in the virtual navigation pre-session. Consistent with our hypothesis, in-depth analyses of grid orientation revealed that grid orientations during the illusion-inducing SYNC condition were similar to those during the navigation pre-session (Supplementary Fig. 6). Of note, our analysis revealed that, unlike the SYNC condition, grid orientation differences between the ASYNC and the virtual navigation pre-session were not statistically clustered (possibly by either instability of the grid code or insufficient magnitude of self-location drift for a robust estimation of grid orientation in the ASYNC condition). Additional analyses also showed that grid orientations during the visuo-tactile stimulation were similar to grid orientations during the self-location report (i.e., active virtual navigation immediately following the visuo-tactile stimulation phase; Fig. 4a). Further corroborating this link, GCLR during the visuo-tactile stimulation phase was correlated with GCLR during the self-location report phase (Fig. 4b), whose spatial vector was matched to the illusory drift by design. Critically, this correlation was only observed for larger drifts in self-location and was absent for drifts below 0.5 vm. This finding also excludes that the correlation was merely derived from the temporally intermingled fMRI dataset of the two phases (i.e., visuo-tactile stimulation and self-location reports). Altogether, the present data suggest that the two types of self-location changes in the current study – (1) illusory drift in bodily self-location elicited by synchronous visuo-tactile stimulation and (2) virtual navigation associated with changes in first-person visual perspective and landmarks in the same virtual room – were represented by a similar grid code and arguably based on the same cognitive map^42, 43^. Changes in the grid orientation of grid cells have been related to hippocampal remapping (more strictly, global remapping; defined as a total reset of hippocampal place fields to encode distinct contexts) and this has been argued to indicate differences in cognitive maps and spatial processes encoding spatial information in different contexts^42, 44, 45^. Accordingly, the present grid orientation data suggest that similar cognitive maps are used during virtual navigation and illusory drifts in self-location, relying primarily on visual environmental signals versus visuo-tactile bodily signals, respectively. It could be argued that the present GCLR findings, during SYNC visuo-tactile stimulation, are related to entorhinal GCLR previously observed during imagined or simulated virtual navigation^28, 29^. However, we do not think this is very likely for several reasons. First, the paradigms (imagination-simulation versus multisensory perception) strongly differ. Imagined navigation or mental simulation of navigation are active cognitive processes where participants were explicitly instructed to imagine or simulate navigation. In the present study, participants were passively exposed to visuo-tactile stimulation and received no instruction to imagine or simulate navigation. Second, we note that GCLR was only observed in the SYNC condition, which was matched in all aspects to the ASYNC condition except for the timing of visuo-tactile stroking. If the present GCLR was based on covert imagined navigation (during visuo-tactile stimulation) this should have also occurred during the asynchronous stimulation, which was not the case. Finally, we observed that the amplitude of GCLR during the illusion-inducing visuo-tactile stimulation period was proportional to the magnitude of illusory forward drifts in bodily self-location within the virtual room, again making it unlikely that the present GCLR signals are based on mechanisms of imagined navigation. Thus, whereas the present SYNC-specific, direction-specific, and drift magnitude-specific effects are typical in multisensory own body perception^12–15^, they have never been reported in imagined navigation. The present method and data also show that it is possible to overcome the disconnect between a participant’s mostly immobile body ‘in’ the MRI scanner and her/his virtual navigation ‘in’ the visual environment^11^. Thus, multisensory stimulation does not only allow to induce illusory drifts in self-location and the projection of the participant towards the avatar and into the virtual environment, but will allow to manipulate other important multisensory aspects of own body perception and their role in navigation and GCLR. This could include, beyond visuo-tactile delay as tested here, different avatar distances, avatar orientations^15, 16, 46^, as well as social aspects of one’s own body and the avatar (e.g., differences in age, gender, or race)^47–50^.

## Methods

### Preregistration

The experimental design and main analyses of this study were preregistered (https://osf.io/7hejk/).

### Participants

Thirty-three healthy participants (20 females; mean age 25.1±1.42) with normal or corrected-to-normal vision were recruited through the online recruitment system open to the general population. The number of participants followed the preregistration and 10% of possible drop-outs were considered (30 + 3). All participants were unaware of the purpose of the research and have no history of psychological disorders. The participants provided informed consent in accordance with the institutional ethical guidelines (IRB #: GE 15-273) and the Declaration of Helsinki (2013). One male participant who kept falling asleep in the scanner was excluded from the entire data analyses, and a total of thirty-two participants were included in the reported results. Three sessions from three participants (one session each) were excluded from the behavioral analyses because of the high variability in the reported self-locations within a session (the variabilities were above 3 standard deviations from the mean).

### MRI-compatible Virtual Reality (VR) and motion capture system

The MRI-compatible visual system (HD system, NordicNeuroLab) is composed of two full high definition (HD) resolution displays (1920 x 1200, 16:10, WUXGA) rendering in 3D/stereoscopic at 60Hz. It provides a diagonal field of view of 60° (52.1° horizontal, 34° vertical). It is also equipped with a built-in diopter correction (−7 to +5) and pupil distance adjustment (44 to 75mm) that facilitates the installation for each participant.

We use a Qualisys MRI-compatible motion capture system composed of six Qualisys Oqus 500+m cameras (recording video at 180Hz with a resolution of 4 Mega Pixel) to track, with the help of reflective markers, the movements of a touch stick used to apply tactile stimuli on the participants’ trunk. The movements of the touch stick were coupled with those of a virtual stick in VR (touching the corresponding body part of the virtual avatar) depending on the experimental conditions. The motion capture system was also used to track the exact location of a participant in the scanner used to simulate the virtual scene they saw during exit and entrance, accordingly (see below for the detail).

### Virtual Scene Simulation for MRI entrance and exit

In order to smoothen the transition between the real and virtual world, we used a virtual MRI room that was an exact 3D model of the actual MRI room. Once the participants were set on the MRI table with the Head Mounted Display, they saw a first-person view from a supine posture looking up at the ceiling of the virtual room. During the entrance to the MRI coils, the first-person view progressively shifted inside according to their exact location captured by the motion capture markers. Coupled with the location of the first-person view, its angle gradually rotates to look in the horizontal direction. The scene was simulated inversely while they were exiting from the scanner. This transfer scene was used to enhance the spatial immersion into the VR environment throughout the entire VR paradigm. The virtual MRI room was only used at the beginning and the end of the experiment. For the main task, participants were transferred to another virtual room with the VR environment for the main experiment (see Fig. 1b)

### MRI-compatible VR: virtual environment and main task flow

The VR paradigm including the virtual room was implemented with Unity Engine (Unity Technologies). The virtual room has a dimension of 12 x 12 virtual meter (vm; matched to meter in real world), containing landmarks (e.g. windows, drawers, couch, painting) only along its walls to avoid a restriction of the virtual navigation or the placement of the avatar while still providing environmental cues (Fig. 1b).

The main task consisted of four FBI-induction sessions, during which participants were exposed to the visuo-tactile stimulation (Fig 1b), and four virtual navigation pre-sessions, during which participants performed the virtual navigation tasks (Fig. 1e). FBI-induction sessions and virtual navigation pre-sessions were given in alternating order, while a virtual navigation pre-session always preceded a FBI-induction session (Fig. 1d). During the experiment, participants could navigate in the virtual room, when required by the task, and autonomously answer questions by using an MRI-compatible button box (Fiber Optic Response Pad, Current Designs) and following the instruction displayed on the VR screen.

### MRI-compatible VR: Virtual navigation pre-session

The virtual navigation pre-session was designed to acquire conventional active navigation and GCLR data during it. The pre-sessions were composed of two kinds of phases: target reaching and target-location report (Fig. 1e; Supplementary Video 1). During target reaching trials, participants actively navigated to reach a target indicated by a light pillar in the virtual room. The target was set at one of the 18 predefined locations 2.5 vm away from the center of the room and spanning around the room every 20°. These locations matched the position where the virtual avatar was placed during the stimulation phase (see Methods for FBI-induction session; Fig. 1). During each trial, participants started at the center of the room and the target was always visible in the first scene they were looking at. Each pre-session had 36 trials of target reaching.

In nine target reaching trials out of 36 (randomly selected), our participants performed target-location report (Fig. 1e): they were instructed to report the last target location (i.e., where they arrived at the end of the preceding target reaching trial). During the target-location report phase, participants’ location in the virtual room was displaced to a position located behind the center of the virtual room (by a variable distance of 2∼3 vm, at random). Then, they actively navigated to the place to report (i.e., the last target location) while the direction of navigation was fixed towards the correct target point (i.e., the expected response). This was designed to assure that the virtual navigation traces during the period of target-location reporting (i.e., a series of short, discrete, and heading-direction-fixed virtual navigations in the virtual room) were sufficient to generate detectable GCLR, which were expected to be similar to the series of self-location drifts induced during the FBI-induction sessions.

### MRI-compatible VR: FBI-induction session

The FBI-induction sessions were designed to (1) induce changes in illusory perceived self-location (drift towards the avatar) that depended on visuo-tactile stimulation condition (SYNC or ASYNC) and (2) to detect potential GCLR and its changes across stimulation conditions. During the visuo-tactile stimulation phase of the session, participants were placed at the center of the virtual room and instructed to look at a virtual avatar placed 2.5 vm in front of them (Fig.1). The value of 2.5 vm was chosen based on the literature on the induction of the Full-Body Illusion^22^, and it was adjusted considering the supine posture of the avatar to ensure an adequate drift in self-location. The avatar was displayed lying down on a virtual table to match the participant’s position in the scanner. The avatar was always seen at the same distance and from the top of the head (to control visual components between the conditions), but its position changed across trials across 18 different angles, spanning the entire 360° of the virtual room (i.e., every 20°). Thus, visuo-tactile stimulations were performed 18 times during a single stimulation session for each of the 20° directions in randomized order (Fig.1b). Synchronous visuo-tactile stimulation lasted for 30 seconds, during which the participant’s abdomen was stroked (and online tracked) while the virtual avatar was being stroked on the abdomen by a virtual stick matching the movement of the real one (Fig.1a; SYNC stimulation; Supplementary Video 2). In the ASYNC condition, the movement of the virtual stick was delayed by 1 second and additionally right-left inverted (Supplementary Video 3). Immediately following each stimulation phase, we asked participants to report their perceived self-location by actively navigating to the position in the virtual room where they perceived as located during the stimulation phase (self-location report phase of Fig.1a). During this phase, there was no virtual avatar shown in the virtual room and participants always started from a position behind the center of the room (by a distance of 2∼3 vm, at random), while keeping the direction they had during the preceding stimulation phase (i.e., same 20° direction). The instruction of the self-location report (i.e., “Go back to where you were”, Fig. 1g) was designed to be ambiguous, not specifying whether the target is the avatar or the first-persion viewpoint during the preceding stimulation phase. The self-location report of a stimulation session consisted of 18 report trials. For each condition (SYNC and ASYNC), two stimulation sessions were performed.

With this self-location report procedure, we not only acquired a measure of the perceived self-location but also had the possibility to determine whether any GCLR is generated by the active virtual navigation during the FBI-induction session (in addition to the virtual navigation pre-session).

Of note, the navigation vectors (i.e., direction, distance) during virtual navigation pre-session were designed to be comparable to the predicted spatial vectors of illusory drift experienced during the visuo-tactile stimulation (i.e., illusory self-location changes; see Fig.1c & f). This was obtained, by using the same 18 locations across the room for both the targets during virtual navigation pre-sessions, and the avatar during FBI sessions. The number of discrete navigations (or self-location changes) during the self-location report phase during FBI induction session was also matched to the target-location report during virtual navigation pre-session (36 times per participant). In addition, the reported target locations during the virtual navigation pre-sessions (which had a correct location to report, unlike the subjective self-location report after the visuo-tactile stimulation) enabled us to assess the reliability of participants’ self-location reports across the entire experiment (Supplementary Fig. 2, also see the prescreening and training section below).

### Questionnaire

Questionnaires regarding bodily self-consciousness were administered after completing the main MRI paradigm. For each condition (SYNC or ASYNC), the visuo-tactile stimulation phase was repeated (in counter-balanced order), followed by five questions shown on the virtual screen. Participants answered the questions by using the button box, rating their level of agreement on a seven-point Likert scale, from 0 (Not at all) to 6 (Very strongly), based on the experience of the preceding stimulation phase. The first question (Q1 - “I felt as if the touch I felt was located where I saw the stroking”) was designed to measure the experience of illusory touch. The second question (Q2 - “I felt as if the body I saw was me”) was to quantify the illusory self-identification to the avatar. The third question (Q3 - “I felt as if I was drifting forward”) was to probe any subjective feeling of the self-location drift toward the avatar. The fourth question (Q4 - “I felt as if I was located in the virtual environment”) reflected presence (i.e. subjective experience of being located in VR) of participants in the virtual environment^51, 52^. The fifth question (Q5 - “I felt as if I had 3 bodies”) was a control question to assess a suggestibility effect on the questionnaire results. A brief debriefing followed once participants completed the experiment.

### Prescreening and training of the participants

The stereoscopic vision of participants was tested before the experiment in the scanner to ensure each of them could fully utilize the spatial information from the MRI-compatible VR goggles. Then, the participants were trained to perform the virtual navigation pre-session outside of the MRI scanner. Unlike in the target-location report phase during the virtual navigation pre-session in the scanner, during training they received feedback on how far from the correct location their reported location was. They kept performing training trials until their performance reached a pre-defined threshold level (mean distance error < 0.5 vm).

Before starting the main task in the MRI scanner, participants went through a familiarization procedure inside the scanner that lasted around ten minutes (without scanning; Fig. 1d). During familiarization, participants were instructed to look carefully at and memorize the virtual room in which they would perform the VR tasks. Then, participants were exposed to the SYNC and ASYNC stimulations (3 trials per condition) and to the following self-Location report phases. These pre-exposures were implemented to minimize possible order effects as well as to train the participants on how to go through the FBI paradigm properly in the main task.

### MRI data acquisition

MRI data were acquired with a 3T MRI scanner (SIEMENS, MAGNETOM Prisma) at the Human Neuroscience Platform of the Campus Biotech (Geneva, Switzerland). The structural images were collected both with a T1-weighted MPRAGE sequence (1 mm isotropic voxels, Number of slices = 208, TR = 2300 ms, TE = 2.25 ms, TI = 900ms) and with a T2-weighted sequence (0.8 mm isotropic voxels, Number of slices = 208, TR = 3000 ms, TE = 409 ms). A T2*-weighted Echo Planar Imaging (EPI) sequence (2 mm isotropic voxels, TR = 1500 ms, TE = 30 ms, Number of slices = 69, Multiband factor = 3, GRAPPA = 2, FoV = 224 mm) was used to acquire functional images covering the entire cerebrum. Both magnitude and phase maps of the B0 field were acquired to correct EPI distortion.

### Regions of interest (ROI) definition for EC

EC ROI of each participant was defined using Freesurfer (v6.0.0) following previous studies^7, 16^. The cortical parcellation was performed with a T1-weighted structural image based on the Destrieux Atlas^53^, referring also to a T2-weighted image. The bilateral EC labels created from the parcellation were examined manually by overlapping them on the corresponding T1- and T2-weighted images. The manual examination was based on anatomical landmarks as described in the previous literature^54–56^. Finally, the examined 3D volume EC ROIs were coregistered and resliced to the mean EPI image, together with the T1-weighted structural image^7, 57^.

### fMRI data preprocessing

The functional images were preprocessed using SPM12. The images were corrected for slice acquisition time, head movement, and inhomogeneous B0 field map. Then, the images were co-registered with the T1-weighted structural image and smoothed with a 5mm full-width-half-maximum Gaussian smoothing kernel. Following previous studies, GCLR analyses (detailed below) were conducted in the native space of each participant without normalization to avoid additional signal distortion^7, 16^.

### Analysis of conventional grid-cell like representation (GCLR) for the Virtual navigation pre-sessions

The Grid Code Analysis Toolbox (GridCAT v1.03) based on SPM12 was used to analyze GCLR from the fMRI data^33^. The procedure consisted of two major parts. In the first part, a putative grid orientation was calculated from a subset of the data. The first generalized linear model (GLM) on the fMRI data estimated *β_cos_* and *β_sin_*, using two parametric modulation regressors: *cos*(6*θ*) and *sin*(6*θ*) based on heading direction(*θ*). Sequentially, voxel-wise amplitude(*A*) and grid orientation(*φ*) were calculated with the estimated betas:

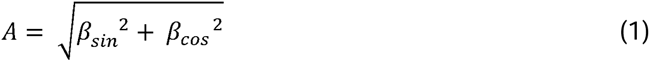

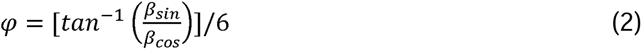

Finally, to determine the putative grid orientation in the EC ROI, the grid orientation(*φ*) of every voxel in the ROI was averaged using the voxel-wise amplitude (*A*) as a weight for each voxel. In the second part, the amplitude of hexadirectional modulation (i.e., GCLR) was calculated from the remaining subset of data (mutually exclusive to the subset of the first part). The hexadirectional modulation was calculated based on the determined grid orientation(*φ*) by contrasting regressors for navigation toward grid-aligned directions (*θ*_*aligned*_) vs. grid-misaligned directions (*θ*_*misaligned*_), where (*θ* − *φ*) *mociulo* 60° = 0° or 30° respectively.

In our case, the procedure of data splitting and estimation of the hexadirectional modulation in segregated subsets were based on a session-wise leave-one-out-cross-validation. Thus, the GCRL of each virtual navigation pre-session was calculated using the grid orientation estimated from the other three pre-sessions. In detail, *β*_*cos*_ and *β*_*sin*_, were estimated per session and averaged across the three sessions. Then, a single grid orientation was estimated through the arctan computation using the pair of the averaged betas. This procedure allowed for the robust estimation of GCLR through repeated calculations while avoiding double-dipping. Finally, the four session-wise values of GCLR were averaged to quantify the magnitude of GCLR for each participant. Of note, we calculate GCLRs during the target reaching and target-location report phases separately (Supplementary Fig. 4), using distinct GLM procedures (e.g., sine and cosine regressors) as described above.

### Calculation of GCLR during the illusion-induction sessions based on the grid orientation of the Virtual navigation control

GCLR during each phase of an FBI-induction session was calculated using the putative grid orientation estimated from the four virtual navigation pre-sessions considered all together (i.e., specifically, target-location report phases during the four pre-sessions; Fig. 1e and Supplementary Fig. 4), on the premise that GCLR induced by illusory self-location changes would be comparable to the conventional GCLR during active virtual navigation with matched self-location changes. As described above, the hexadirectional modulation during either SYNC or ASYNC stimulation phase was calculated by contrasting regressors for the stimulation in the grid-aligned directions vs. grid-misaligned directions. Finally, using the same method, we also calculated the GCLR during the self-location report phase in the FBI-induction session, where our participants actively navigated to report their perceived self-location.

### Mediation Analysis

Mediation analysis (Supplementary Fig. 7b) was performed with a dedicated R package (mediation, v4.5.0). For the mediation analysis, round-wise data with the full range of categorical (Condition; SYNC or ASYNC) and numerical variables (reported self-location, GCLR) were used. The simulation count was set to 3000..

### Statistics and Reproducibility

R (v3.6.3 for Windows) and RStudio (v1.1.423) were used for the statistical analyses of the results. The existence of GCLR was assessed with a non-parametric one-sided Wilcoxon signed-rank test based on the a priori hypothesis that its mean was greater than 0. Any comparison between amplitudes of different GCLRs was performed with a two-sided Wilcoxon signed-rank test. Questionnaire ratings were also compared with a two-sided Wilcoxon signed-rank test. For the reported self-location data, outlier trials outside of the three standard deviation range and miss trials without any button manipulation prior to answer were excluded from the statistical analysis (0.51%; 18 out of 3564). To maximally utilize the data points, the R package for mixed-effects regressions (lme4, v1.1.26) was used to statistically assess the condition-wise difference (Supplementary Table 1). Correlations between parameters were assessed by the mixed-effects models. Exceptionally, the correlation between GCLRs during visuo-tactile stimulation phase and self-location report phase (Fig. 4b) was assessed with a Spearman correlation because the data points were not enough for the mixed effects models.Statistical analyses and visualization of the circular data (e.g., grid orientations) were performed using the MATLAB toolboxes (CircStat, CircHist). First, directionality of the circular distribution was assessed by the Rayleigh’s test for non-uniformity. Once the distribution was determined as non-uniform, its circular mean, confidence intervals, and r value (i.e., resultant vector length) were calculated.

## Acknowledgments

This work was supported by the Korean Government Scholarship Program for study overseas, the Korea Institute of Science and Technology (KIST) Institutional Program (2E31642), and the Bertarelli Foundation to H.-J.M. O.B. is supported by the Swiss National Science Foundation (No. 320030_188798) and by the Bertarelli Foundation. Additional support was provided by the Fondation Campus Biotech Geneva (FCBG)—a foundation of the Swiss Federal Institute of Technology Lausanne (EPFL), the University of Geneva (UniGe), and the Hôpitaux Universitaires de Genève (HUG), the Institute of Translational Molecular Imaging (ITMI).

## Author Contributions

Conceptualization, H.-J.M., H.-D.P., B.G., C.T., and O.B.; Methodology, H.-J.M. and L.A.; Software, H.-J.M. and L.A.; Formal Analysis, H.-J.M.; Investigation, H.-J.M., L.A., and C.T.; Writing – Original Draft, H.-J.M.; Writing – Review & Editing, E.D.F., B.G., H.-D.P., and O.B.; Resources, H.-J.M. and L.A.; Funding Acquisition, O.B.; Supervision, O.B.

## Data and code availability

The data that support the findings of this study and the analysis code are available from open respository (https://osf.io/7hejk/).

