## Supplementary Information for "Changes in spatial self-consciousness elicit grid cell-like representation in entorhinal cortex"

**Supplementary Text**

**Previous Bodily Self-Consciousness (BSC) studies investigating self-location changes and related brain activity.**

Drifts in self-location have been induced previously using multisensory stimulation, but never in virtual navigation settings during fMRI as in the present study. Previous work applied visuo-tactile stimulation either to the back of standing participants and to the back of an avatar^1-3^ or to the back of supine participants and the back of an avatar (adaptation to fMRI) and found to activate angular and posterior temporal gyrus^4^. Other bodily illusion research applied visuo-tactile stimulation to the abdomen in supine participants and an avatar during fMRI^5-7^, reporting recruitment of a network consisting of retrosplenial cortex, intraparietal sulcus, posterior cingulate cortex, and hippocampus. However, Ionta et al. (2011)^4^ elicited drifts in the vertical direction, while latter studies did not quantify drifts in self-location, and drifts were never investigated across many angles in virtual rooms and never with respect to activity in EC nor the GCLR, in particular.

**Reliable measure of experienced self-location: through pre-training and the correction based on the target-location report**

To ensure the reliability and precision of the self-location reports, we pre-trained our participants before the scan so that they learned to accurately report a location in the virtual room (see Methods). Also, since participants performed the target-location report during the virtual navigation pre-session preceding FBI-induction (where they had to report their location at the end of ‘standard’ virtual navigation), we could use those as a baseline of the precision/bias of their reports. Based on the control analysis, we further confirmed a forward drift during the induction of the illusion even when correcting for the subject-wise biases in the report (e.g., some participants may have a bias to always report themselves closer to the center of the room, which would affect their self-location report after the visuo-tactile stimulation; Supplementary Figure 2c-d). Hence, our behavioral results confirmed and extended the findings from previous FBI studies, providing self-location drifts on the horizontal plane and its quantifiable measures.

**Influence of an initial position on the self-location report**

**At the beginning of each self-location report phase, our participants started from a position of 2~3 vm behind the center of the room (distances were pseudo-randomized). We assessed whether the difference in initial position affected the reported self-location through the dedicated mixed-effects model: SL_report ~ Initial_distance + Condition + (1|Subject/HeadDirection). Data show that the initial distance from the center of the room influenced participants’ self-location report (df = 1, F = 10.152, p = 1.47e-03; see figure below): the farther their initial location was away from the avatar, the farther their reported self-location was also away from it. Critically, however, regardless of the effect of the initial point, the effect of the stimulation condition (SYNC vs. ASYNC) on the reported self-location remained significant (F = 44.196; p = 4.04e-11). In addition, the GCLR was proportional to the reported self-location (df = 1, F = 4.44, p = 0.042). Notably, the initial location cannot account for the difference in GCLR, because GCLR was estimated from the session-wise data, where initial locations were equally pseudo-randomized. Finally, the navigated distance also was not correlated with the GCLR (df = 1, F = 0.01, p = 0.92; Supplementary Figure 8). Critically, we underscore that the two visuo-tactile conditions were matched in terms of angles and distances from the avatar (while their orders were randomized), excluding that these parameters can account for the differences in behavior and GCLR between the conditions.**

**Absence of GCLR during the target reaching phase of the virtual navigation pre-session**

**In the virtual navigation pre-session, GCLR was significant during the target-location report phase, but not during the target reaching phase, although participants performed active virtual navigation during both phases (Supplementary Figure 4). The absence of GCLR during the target reaching could indicate that the grid-like code was not or less recruited during the target reaching trials. Indeed, it has been suggested that the hippocampal-entorhinal structures are not strongly recruited when an allocentric spatial strategy is not necessary to recognize/update self-location (e.g., during beacon-guided navigation;^8-10^). During a target reaching trial, participants had to reach a target seen right in front of their viewpoint. They did not need to locate themselves in the room to do the task (and only their egocentric distance to the seen target object should be updated). They had to simply memorize where the target was located in the room in order to perform the following target-location report phase (i.e., no need to recognize/update allocentric ‘self’-location). Therefore, given these details, it is reasonable that GCLR was detected during the target-location report phase but not during the target reaching phase.**

**One might argue that more active processes with memory recall component during the target-location report phase (that was absent during the target reach phase) might have led to the observed difference in GCLR. However, during the stimulation (FBI) phase, participants were ‘passively’ stimulated, but the presence of GCLR related to the illusory self-location drift indicates that the presence of an ‘active’ process is not necessary to elicit GCLR. This is further supported by GCLR during ‘passive’ observation of someone else’s navigation ^11^.**

**We also note that target reaching trials were very short (~4.83s and net navigation ~1.55 s on average) and densely packed with short inter-stimulus intervals (~1.2 s), which is not ideal for estimating GCLR. This might have contributed to insignificant GCLR in the target reaching phase. However, the target-location report phase, where very robust GCLR was observed (Supplementary Figure 4), was comparably short (~ 6.13s and net navigation ~3.12 s on average) and had only a quarter of trials per session than the target reaching phase (9 vs. 36). Self-location report phase also had a comparable mean duration (~ 5.62 s and net navigation ~2.22 s; 18 trials per session). Based on these observations, we believe it is very unlikely that the absence of GCLR was merely due to the poor data quality or lack of the statistical power.**

**GCLR estimation exclusively using data from the FBI-induction phase**

We could not detect significant GCLR by exclusively using the data of the FBI-induction phase (SYNC/ASYNC). Given that the GCLR estimation procedures require splitting the dataset into, at least, two subsets (see Methods), this was possibly due to the lower statistical power of the FBI-induction sessions where we had only two sessions per kind of stimulation (SYNC and ASYNC) compared to the four virtual navigation pre-sessions. Additionally, the signal-to-noise ratio could have been lower in the GCLR during the stimulation phase than the virtual navigation, as suggested by the weaker voxel-wise six-fold modulation (Supplementary Fig. 4b).

**GCLR proportional to the illusory drift but not to navigation distance during virtual navigation**

In the previous reports^12, 13^, human GCLR during navigation has never been related to the navigation distance. Congruently, we did not observe a significant correlation between navigation distance and GCLR during active virtual navigation (i.e., virtual navigation pre-session or self-location report phase; Supplementary Figure 7). Then, why did the changes in the self-location correlate with the GCLR during illusion-inducing multisensory stimulation? A possible explanation is that the drift in self-location reported by the participants may represent the strength or vividness of the subjective experience of forward drift (distinct from the distance of the drift, per se), as reflected in the significant correlation between reported drift and questionnaire item (Fig. 2b). Hence, the stronger experience of illusory self-location change might also contribute to the difference in GCLR between SYNC and ASYNC. As an alternative explanation, a minimum spatial displacement might be necessary to generate detectable GCLR through fMRI analysis. This would imply that on trials when the drift was too small, the measured GCLR would not be detectable (i.e., close to zero) and this would generate the correlation observed. It might also explain the non-significant GCLR in ASYNC, which had smaller (thus, more likely sub-threshold) drifts than SYNC. From the single-unit study of human grid cells, it has been suggested that human grid cells have spacing between their grids of 1 to 6 m^14, 15^). Therefore, considering that the radius of each firing field is usually around one-third of the grid spacing^16^, the minimum displacement necessary to induce detectable changes in firing rates of grid cells should be between 0.3 and 2 vm, which would be consistent with what we observed (~0.5 vm; Fig. 4b & Supplementary Fig. 6a).

One might argue that fMRI-based GCLR (i.e., the characterized modulation of BOLD activities by heading directions), which presumably originated from conjunctive grid cells (i.e., head direction + grid cells)^12^, should be maintained even with the minimal movements (regardless of the size of the firing field). However, in addition to the directional selectivity, fMRI-based GCLR requires changes in self-location (as conjunctive GCs are not just head direction cells; they fire more when an agent is ‘passing’ the spot in a preferred direction). In support of this, no movements or slow movements do not generate GCLR as reported in Doeller et al. (2010)^12^ and in Horner et al. (2016)^17^. Accordingly, we argue that there likely is a threshold level of self-location changes to evoke fMRI-based GCLR. Further investigations are required on these questions.

**Changes in voxel-wise sixfold modulation might imply rate-remapping during the visuo-tactile stimulation phase.**

Although the grid orientations were similar and no significant difference in GCLRs (i.e., sixfold modulation calculated in the entire EC) was observed between SYNC stimulation and self-location report phase, the voxel-wise sixfold modulation (or voxel-wise Amplitude($A$), see conventional GCLR analysis section of the Methods^13^) was significantly different between those two conditions (smaller during the SYNC condition compared virtual navigation in self-location report phase) (Supplementary Fig. 4b){Moon, 2022 #8896}{Moon, 2022 #8896}. Such difference might originate from differences in sensory inputs: during virtual navigation compared to the FBI, the brain is exposed to many additional signals, including visual perspective changes and optic flow, as well as movements guiding the virtual navigation that could, overall, increase the related brain activity. Additionally, rate-remapping may also explain the observed change in voxel-wise sixfold modulations, as a way of encoding self-location changes in the same virtual room, but in different non-spatial contexts^18, 19^. Changes in the firing rates of place cells without changes in their place fields (i.e., rate-remapping) are often associated with non-spatial sensory inputs (e.g., odor)^20^ and are suggested to be associated with rate changes in grid cells^18^. Visuo-tactile stimulations (e.g., touches on the abdomen) in our study might evoke similar brain mechanisms involving hippocampal rate remapping and consequential GCLR changes: magnitudes of the voxel-wise hexadirectional modulation changed while grid orientations were preserved, suggesting a mild change of grid code to update non-spatial context and relevant sensory information.

**Potential influence of movement speed on GCLR**

The speed of virtual navigation has been shown to impact GCLR^12^. Although the difference in GCLRs between the SYNC stimulation and self-location report was not significant, the decreased voxel-wise sixfold modulation during the SYNC compared to self-location report phases might be due to differences between in the “mental speed” of the elicited drifts in perceived self-location versus the speed of displacements during virtual navigation. Even if we cannot precisely estimate the instantaneous “mental speed” of drifts in self-location location or even easily compare the drift and navigation speed, it could be assumed that the average speed of drift in self-location was slower than during virtual navigation: the average speed (navigated distance over time) during the self-location report phase was likely faster than the “speed” during SYNC, because the self-location changes were larger (Supplementary Fig. 2 & 7; they were placed behind the center by 2~3 vm before each report) for a shorter duration of time (on average 5.8 s compared to 30 s during SYNC).

**Supplementary Videos**

**Virtual_navigation_Moon_et_al.mp4**

<https://drive.google.com/file/d/1YlPNXFU8kw5ouIA7zxk2udMWlW-q26T8/view?usp=sharing>

**:** task procedures during virtual navigation session (target reaching and self-location report)

**Stimulation_SYNC_Moon_et_al.mp4**

<https://drive.google.com/file/d/1L-urYeC4p_maei3A6VafLAsTJFfOl1Te/view?usp=sharing>

**:** task procedures during a stimulation session of SYNC condition (stimulation and self-location report)

**Stimulation_ASYNC_Moon_et_al.mp4**

<https://drive.google.com/file/d/1O9pIjF1k6nu6EFmrYNMs31uSsz4JIwJl/view?usp=sharing>

**:** task procedures during a stimulation session of ASYNC condition (stimulation and self-location report)

**Supplementary FIGURES**

**Supplementary Figure 1**

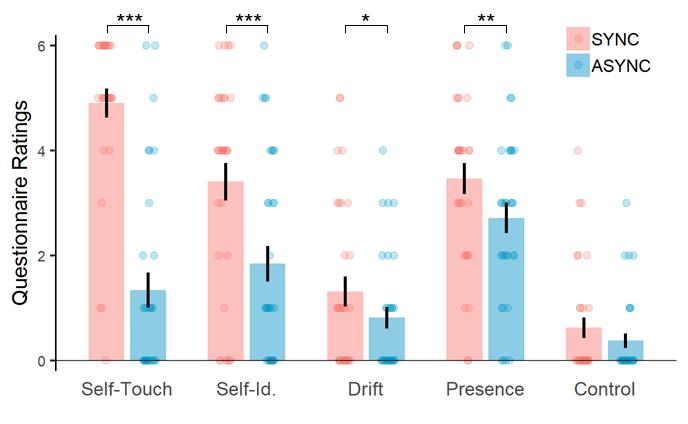

Fig. S1, related to Fig.2, Subjective ratings of the BSC questionnaire

Questionnaires regarding subjective BSC experience were administered in the MRI scanner at the end of the experiment (n = 32). The results showed a significant difference in the illusory ‘Self-Touch’, ‘Self-Identification’, ‘Drift’ in Self-location, and ‘Presence (i.e., feeling of being located in the virtual environment)’ ratings between the two experimental conditions (SYNC vs. ASYNC). We did not find a significant difference between the conditions for the ‘Control’ question. Each error bar indicates a standard error. *: 0.01 <= p < 0.05, ** : 0.001 <= p < 0.01, *** : p < 0.001.

**Supplementary Figure 2**

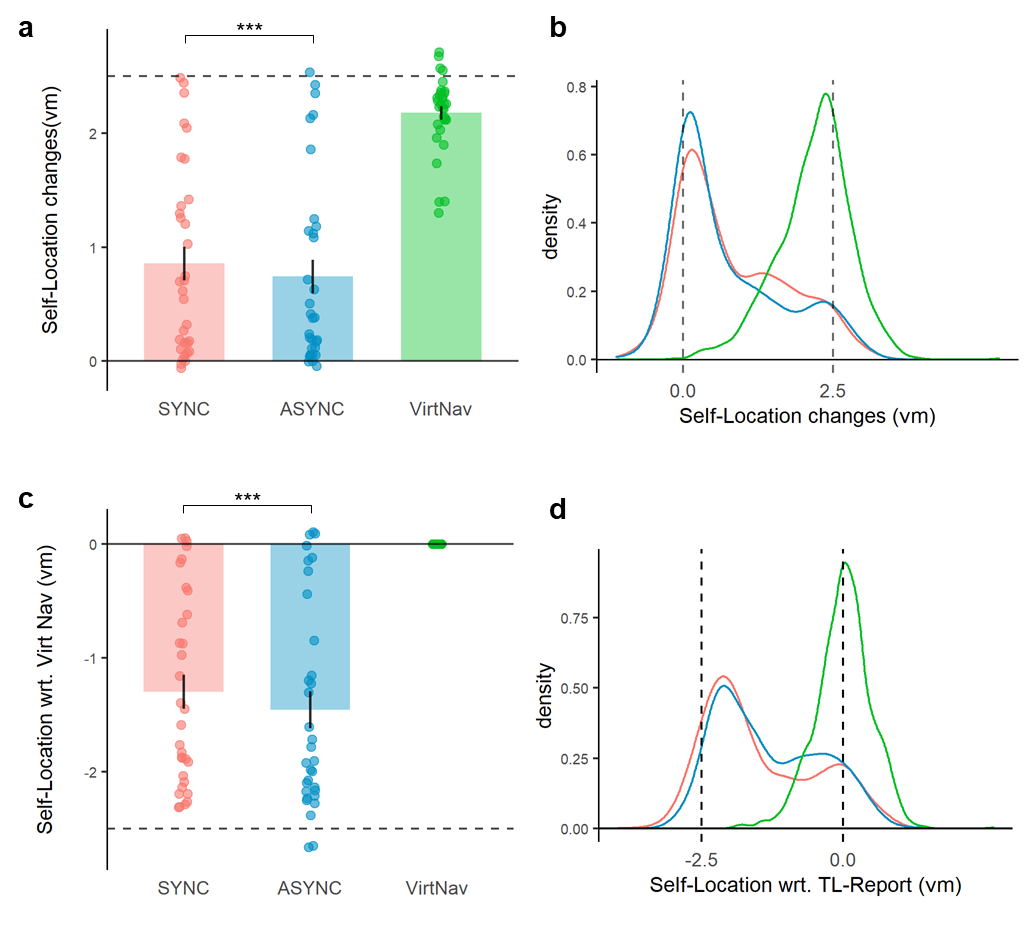

Fig. S2 Self-locations reported after visuo-tactile stimulations and Virtual Navigation tasks

**a,** As shown in Fig. 2a, self-locations reported following the SYNC visuo-tactile stimulation phase were farther away from the viewpoint than the ASYNC stimulation (p < 0.001; n = 32). Of note, self-location reported after the virtual navigation task showed that some participants tend to navigate less far to the target location (They were supposed to navigate to the target at the 2.5 vm point after the virtual navigation task). **b,** A density graph again confirmed the bigger self-location drift in the SYNC than the ASYNC condition. Importantly, the graphs of virtual navigation pre-session (panels a & b) demonstrate the variability of reported self-location across our participants, although, ideally, our participants always had to report self-location changes of 2.5 after the virtual navigation task. This suggests possible differences in the tendency of reporting self-location across the participants: some may report themselves closer to the center than others, which very likely affects self-location reports after the visuo-tactile stimulation. **c,d,** To take into account the possible influence of the participant-wise tendency in the self-location report, we re-calibrated reported self-location changes based on the target-location reports during the preceding virtual navigation pre-session. We subtracted the mean value of the target-location reports during the preceding pre-session (which should be 2.5vm, ideally, but varied possibly reflecting biases) from the self-location-report values of the following FBI-induction session. As a result, we again confirmed the bigger self-location changes in the SYNC condition than the ASYNC condition (p < 0.001, n = 32). Each error bar indicates a standard error. *** : p < 0.001.

**Supplementary Figure 3**

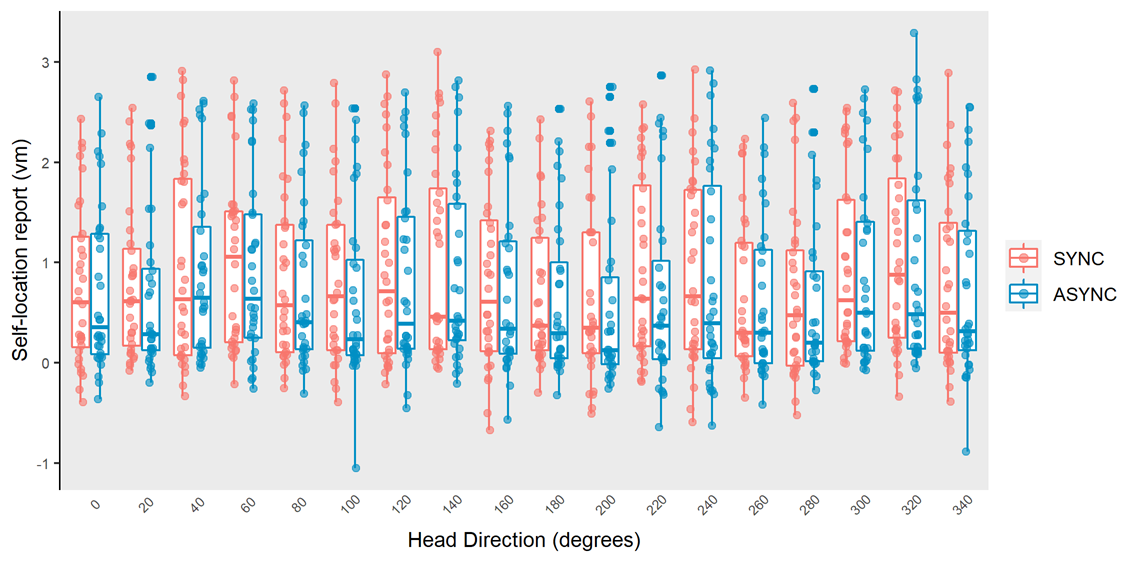

Fig. S3, related to Fig. 2, Reported self-locations by head directions. Possible influence of viewing angle (i.e., head direction in the room) was assessed through Two-way repeated measures ANOVA with head direction and stimulation condition as two within-subject factors. We found that viewing angle did not significantly affect self-location report after the Stimulation phase (Df = 17, F = 0.301, p = 0.997), while the effect of the condition (i.e., SYNC vs. ASYNC) was significant (Df = 1, F = 5.36, p = 0.021). These findings indicate that specific head direction or landmarks in our scene cannot explain the obeserved difference in reported self-location between stimulation conditions.

**Supplementary Figure 4**

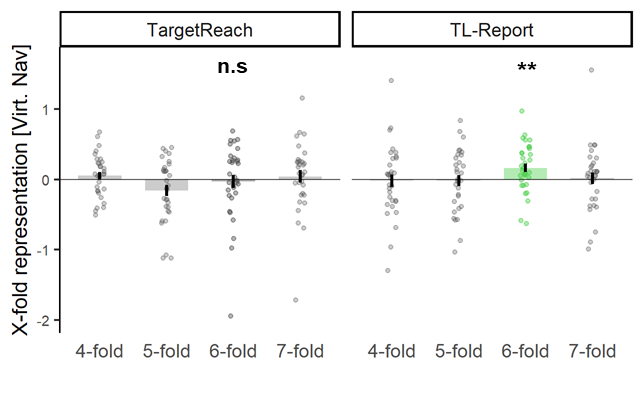

Fig. S4, related to Fig. 3, Multi-fold symmetry BOLD modulation during each phase of Virtual Navigation pre-session. Grid cell-like representation (GCLR) was significant during the target-location report (TL-Report) phase in the Virtual Navigation pre-session, while not during the Target Reaching trials (Fig. 1e). Grid orientation estimated during TL-Report phase of the pre-session was used to estimate GCLR during the FBI-induction session. Each error bar indicates a standard error. n.s.: p > 0.05, ** : 0.001 <= p < 0.01.

**Supplementary Figure 5**

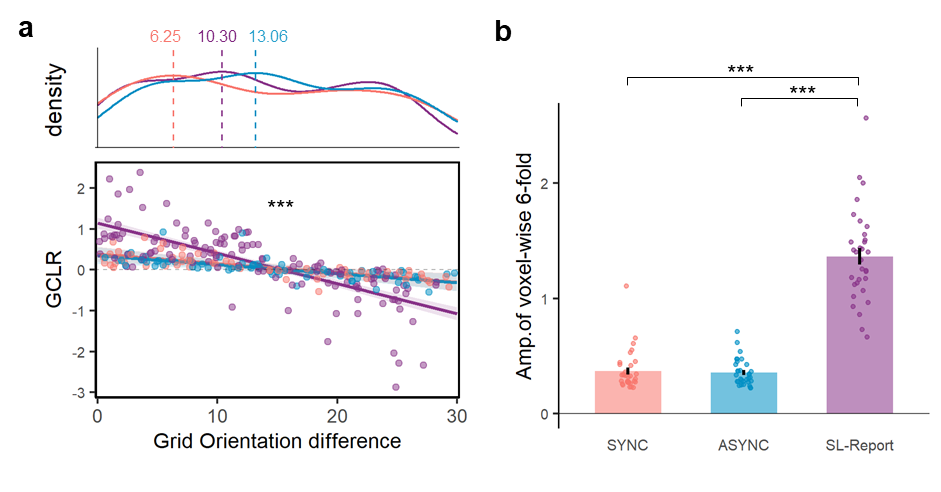

Fig. S5, related to Fig.3, Sub-parameters of GCLRs for different experimental phases during FBI-induction session. a, GCLRs calculated based on the grid orientation during the Virtual navigation pre-session were negatively correlated with the difference of grid orientation during the pre-session and each task phase. For this figure, the grid orientation differences were considered as absolute values, irrespective of the sign (+/-). The dashed line in the density plot indicates the mode value for each phase. b, Voxel-wise hexadirectional modulation was averaged per participant during each task phase. The voxel-wise modulation was calculated per voxel independently of the other voxels in the region of interest (ROI), while GCLR was calculated based on a grid orientation averaged across the entire EC ROI. The results showed that its amplitude was bigger during the self-location report phase than both SYNC and ASYNC visuo-tactile stimulation phases. Each error bar indicates a standard error. n.s.: p > 0.05, *: 0.01 <= p < 0.05, ** : 0.001 <= p < 0.01.

**Supplementary Figure 6**

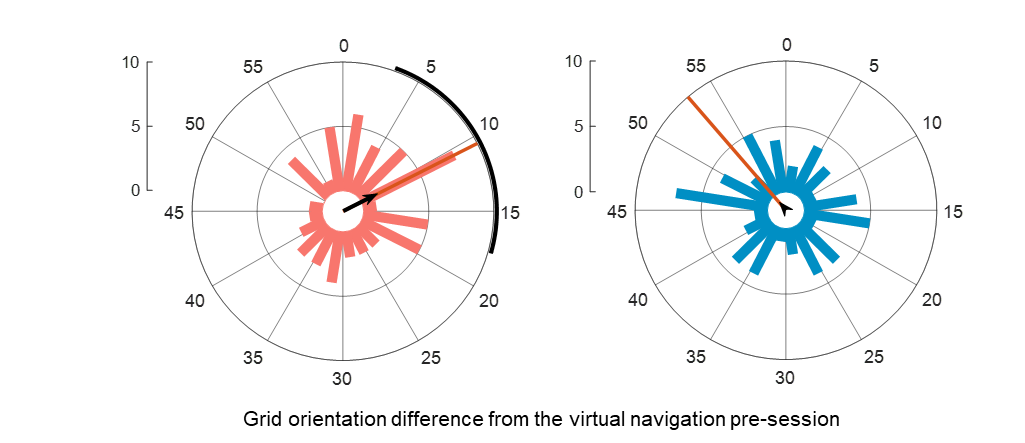

Fig. S6 Difference in grid orientation during the visuo-tactile stimulation and during the virtual navigation pre-session. Grid orientation during the SYNC stimulation phase differed from the virtual navigation pre-session by 10.49° ± 7.11° (r = 0.253). The difference in grid orientation between ASYNC stimulation and the pre-session was 53.24 (r = 0.067). Of note, the difference in ASYNC was not statistically clustered. Thus, the mean difference might not be statistically meaningful and its confidential intervals also could not be estimated. A red line indicates the mean angular difference and the bold black curve at the rim indicates the confidence intervals.

**Supplementary Figure 7**

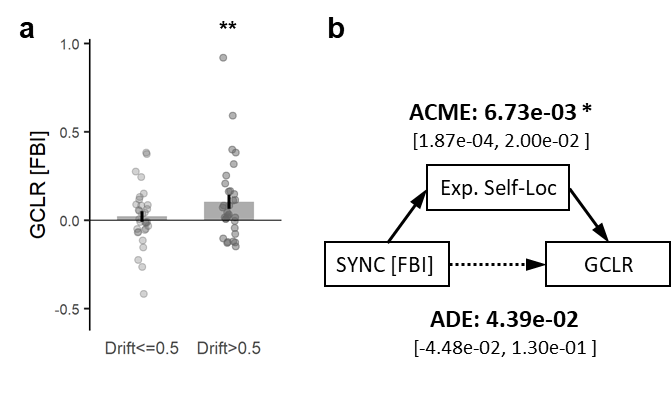

Fig. S7 GCLR during the stimulation phase was related to the drift in the experienced self-location. a, GCLRs during visuo-tactile stimulation were median-split into two groups based on the extent of the forward self-location drift (either Drift > 0.5 or Drift <= 0.5). The results showed that the GCLR was only significant in the bigger drift group (i.e., Drift > 0.5, p = 8.73e-03; Drift <= 0.5, p = 0.244). b, The mediation analysis was performed to further clarify the mechanisms underlying the GCLR during the visuo-tactile stimulation. The analysis suggested that the difference in GCLR between the SYNC and ASYNC conditions was significantly mediated by the difference in the experienced self-location between them (ACME), rather than solely caused by the direct impact of the synchrony of the visuo-tactile stimulation (ADE), per se. For the mediation analysis, round-wise data with the full range of categorical (Condition; SYNC or ASYNC) and numerical variables (reported self-location, GCLR) were used. Each error bar indicates a standard error. ** : 0.001 <= p < 0.01.

**Supplementary Figure 8**

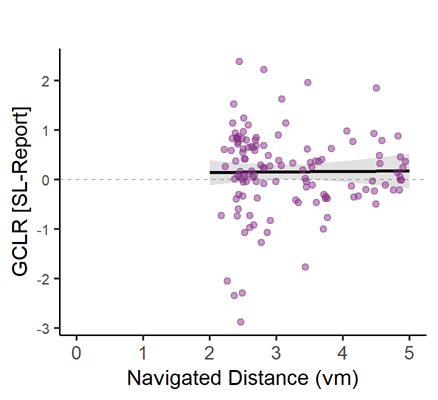

Fig. S8 GCLR and the navigated distance, GCLR during self-location report phase was not significantly correlated to the navigated distance (df = 1, F = 0.01, p = 0.92).

| **Dependent variable** | **Fixed effects** | **Random effects** | **Nested** | **N Subjects** | **N datapoints** |
| --- | --- | --- | --- | --- | --- |
| **Effects of visuo-tactile stimulation and questionnaire ratings on Drifts in Self-location** | | | | | |
| SL-Report | Condition | Subject | HeadDirection | 32 | 2234 |
| SL-Report wrt. TL-Report | Condition | Subject | HeadDirection | 32 | 2234 |
| SL-Report | Condition * Q1 | Subject | HeadDirection | 32 | 2234 |
| SL-Report | Condition * Q2 | Subject | HeadDirection | 32 | 2234 |
| SL-Report | Condition * Q3 | Subject | HeadDirection | 32 | 2234 |
| SL-Report | Condition * Q4 | Subject | HeadDirection | 32 | 2234 |
| **Effects of SL-Report, Navigated distance, and difference in Grid Orientation on GCLR** | | | | | |
| GCLR | SL-Report | Subject | - | 32 | 125 |
| GCLR | Navi-Distance | Subject | - | 32 | 125 |
| GCLR | Condition* Abs_difference_in_G.O. | Subject | - | 32 | 256  (Stimulation  + SL-Report) |

**Supplementary Table**

**Supplementary Table 1.** Model specifications of the used mixed-effects models
